# Viral-mediated fluorescent labeling of hyaluronan reveals extracellular matrix dynamics in the mouse brain in vivo

**DOI:** 10.1101/2025.01.31.635882

**Authors:** Mario Fernández-Ballester, María Ardaya, Nathalie Dutheil, Leslie-Ann Largitte, Abraham Martín, Federico N. Soria

## Abstract

The extracellular matrix (ECM) of the brain is primarily composed of the glycan polymer hyaluronan (HA), a core scaffold that nucleates proteoglycans forming a self-assembled matrix that acts as structural framework and signaling hub. Since most of the neural matrix is composed of sugars, development of genetically encoded tags has been limited. Therefore, although several staining protocols exist for ECM in fixed tissue, there are no reliable matrix labels for live imaging. Here we report a viral-mediated fluorescent probe that binds to HA and labels the mouse brain ECM. The vector encodes the HA binding domain from neurocan fused to GFP and an externalization tag (AAV-Ncan-GFP), enabling transduced cells to secrete the fluorescent hyalectan into the extracellular space, thereby labeling HA. We demonstrate stable probe expression in organotypic brain slices, as well as in vivo in the mouse cortex, where it labels both perineuronal nets and interstitial matrix. We validate HA labeling through colocalization with HABP and sensitivity to hyaluronidase, and confirm the probe’s extracellular localization by shadow imaging. As a proof of concept, we combine AAV-Ncan-GFP with dendritic spine imaging *ex vivo* and calcium transient imaging *in vivo*, providing a real-time map of local ECM alongside neural function. The probe enables time-lapse imaging of ECM dynamics in live mice, facilitating longitudinal studies across a wide range of timescales, from minutes to days. The results establish AAV-Ncan-GFP as a valuable tool for real-time observation of brain ECM and a promising resource to explore ECM dynamics and brain function in vivo.

## INTRODUCTION

The extracellular space (ECS) of the brain contains a dynamic scaffold known as the extracellular matrix (ECM). This dense network of macromolecules, which in the central nervous system (CNS) is secreted by glia and neurons, accounts for 20% of brain volume and occupies the void between neural cells ^1^. Once regarded solely as a physical substrate, increasing evidence supports its role as both neuro- and gliomodulator ^2,3^. The ECM also contributes to brain function by regulating diffusion in the ECS ^4,5^, the central site of neurotransmission.

The brain ECM exhibits distinct configurations ^6^, with the interstitial matrix (iECM) being by far the most prevalent. Unlike the ECM from connective tissue, where collagen is the main unit, the neural interstitial matrix primarily consists on sugar polymers termed glycans ^7^, being hyaluronan (HA) the most abundant of these. Matrix proteoglycans bind to HA forming a gel-like interlaced structure that functions as cell-attaching scaffold when assembled and as a signaling molecule upon disruption ^8,9^. This iECM, thus, fills the extracellular space, surrounding every cell in the brain and serving as critical tissue organizer ^1,4^. However, despite iECM ubiquitous nature, most of our understanding on how the ECM regulates brain function comes from studies on perineuronal nets (PNNs), which are specialized ECM structures predominantly encasing inhibitory interneurons ^10^. This is partly because labeling and manipulating genetically the HA-rich iECM is challenging due to its polysaccharide nature.

Currently, the primary method for visualizing brain ECM involves staining its components such as HA or chondroitin sulphate proteoglycans (CSPG) through binding proteins (*Hyaluronic Acid Binding Protein*, or HABP) ^11^, or lectins (*Wisteria Floribunda Agglutinin*, or WFA staining) ^12^ in fixed tissue. These labeling techniques, combined with enzymatic degradation of ECM components, have been instrumental in demonstrating the role of ECM in functions ranging from synaptic plasticity ^13,14^ to glial physiology ^15–17^. However, because these methods label the ECM post-mortem, they lack the ability to capture its dynamic contribution in real-time. There is an urgent need for techniques that allow simultaneous labeling of the ECM and intravital markers for neurons and glia. Such approaches would enable real-time observation of cell-ECM interaction in a *dynamic* context, providing deeper insights into ECM’s role in brain physiology and pathology.

To address this critical gap, we present a viral-mediated genetically-encoded fluorescent probe that binds to HA and labels the ECM in the mouse brain, a ground-breaking tool for imaging brain ECM dynamics in live brain tissue. The vector encodes the HA-binding domain from neurocan (Ncan) fused to green fluorescent protein (GFP) and a sequence for externalization, labeling HA in the ECS. Originally developed for zebrafish ^18^, we engineered the vector for AAV-mediated stable expression in mice and validated its ability to label brain HA We demonstrate stable probe expression in organotypic brain slices, as well as in vivo in the mouse cortex, where it labels both PNNs and iECM. HA labeling is confirmed through colocalization with HABP and sensitivity to hyaluronidase, with extracellular localization further verified using confocal shadow imaging. As a proof of concept, we combine AAV-Ncan-GFP with dendritic spine and calcium transient imaging *in vivo*, providing a real-time local map of the matrix alongside neural function. Additionally, we perform, for the first time, longitudinal time-lapse imaging of ECM dynamics in live mice over several days.

## RESULTS

### AAV-Ncan-GFP mechanism for iECM labeling

Although several staining protocols exist for brain ECM in fixed tissue, reliable labels for live imaging of the iECM are lacking. This is because HA has a simple, unbranched structure of repeated unsulfated sugars (glucuronic acid and N-acetylglucosamine) that is conserved across species ^19,20^, lacking antigenic structures and, therefore, immunogenicity. Consequently, it is impossible to produce specific antibodies against it ^21^. An additional challenge is that HA, being a sugar and not a protein, cannot be genetically encoded. Therefore, it is not possible to fuse it with a fluorescent protein, as is done with GFP-tagged cell markers for live imaging.

To address hyaluronan vital labeling, we used neurocan-GFP (Ncan-GFP), a fused protein originally conceived to visualize HA in live zebrafish ^18^. Neurocan is a chondroitin sulphate proteoglycan of the lectican family, capable of binding to various ECM components. However, its N-terminal domain has a specific affinity for HA ^22^. Ncan-GFP is composed of only the HA-binding domain of neurocan (Ncan), but fused with GFP. Thus, the probe anchors specifically to HA and labels fluorescently the glycan structure of the brain iECM. We engineered the genetic code for Ncan-GFP into a viral vector, adding regulatory elements to improve its expression in mammals. We then packaged the new plasmid into an adeno-associated virus, isotype 9, (AAV9-Ncan-GFP), that in mice infects primarily neurons and astrocytes. The vector encodes Ncan-GFP with a sequence that directs it to the secretory pathway, allowing transduced cells to secrete the fluorescent binding protein into the ECS, thereby labeling HA, the main component of the iECM (**Figure 1**).

**Fig. 1.**
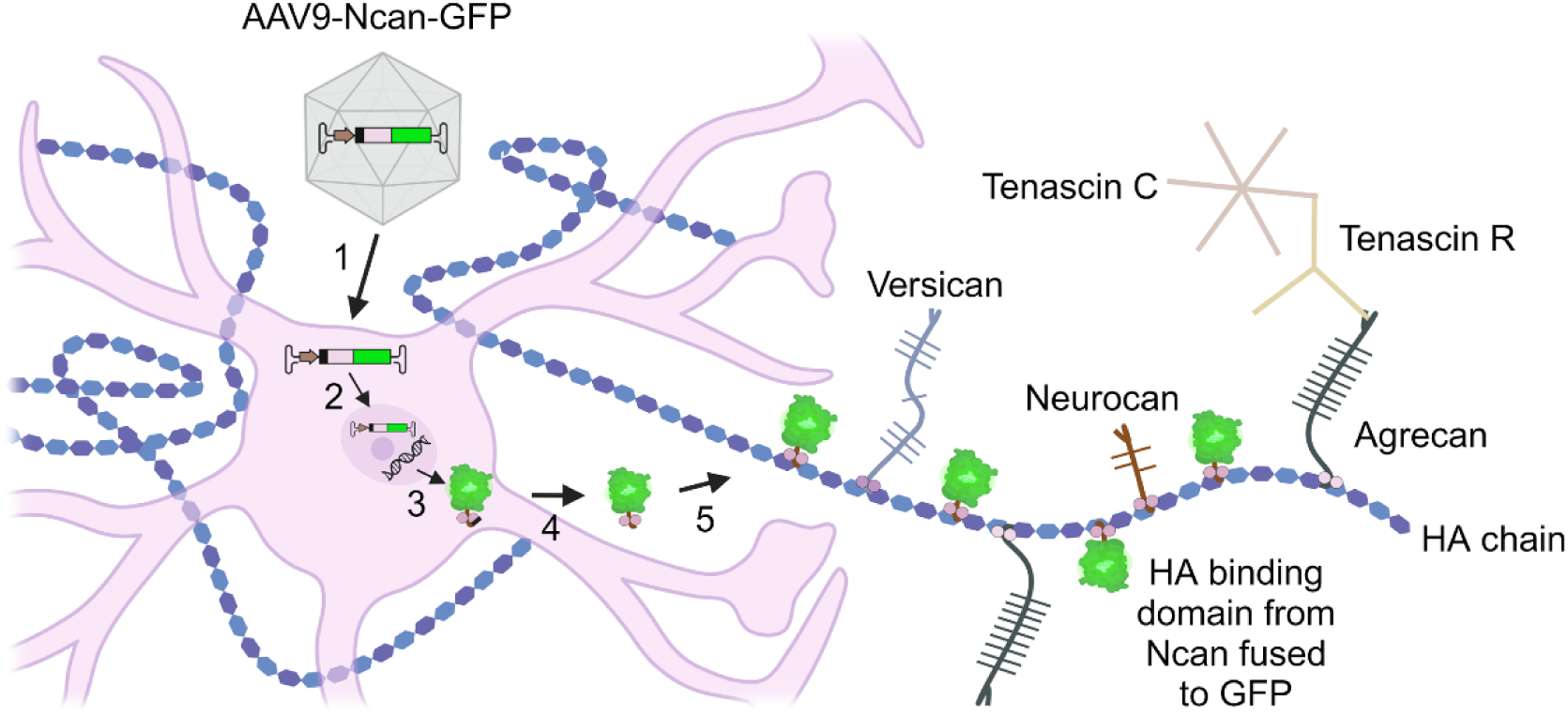
AAV-Neurocan-GFP mechanism for HA labeling. Schematics of AAV-Ncan-GFP cell transduction (1-2), synthesis (3), export to the ECS (4) and binding to hyaluronan (5), thus labeling the brain ECM. This figure was created in BioRender (Academic Individual License).

### AAV-Ncan-GFP labels hyaluronan and reveals iECM heterogeneity

We began by validating HA vital labeling in rat cortico-striatal organotypic culture (OTC) slices. This model preserves tissue architecture and topology of the brain parenchyma, retaining the ECS ^23,24^ and the spatial relationships between brain cells and the ECM ^25,26^. Additionally, it enables long-term genetic and pharmacological manipulation, which can be combined with live imaging at various timescales ^27^. We therefore established a new protocol for OTC and AAV-Ncan-GFP infection ^28^, utilizing the membrane-interface culturing method (**Figure 2a**). Cortico-striatal slices obtained from postnatal day 5-7 rat pups were cultured on membrane inserts and infected with AAV-Ncan-GFP at day-in-vitro (DIV) 3.

**Fig. 2.**
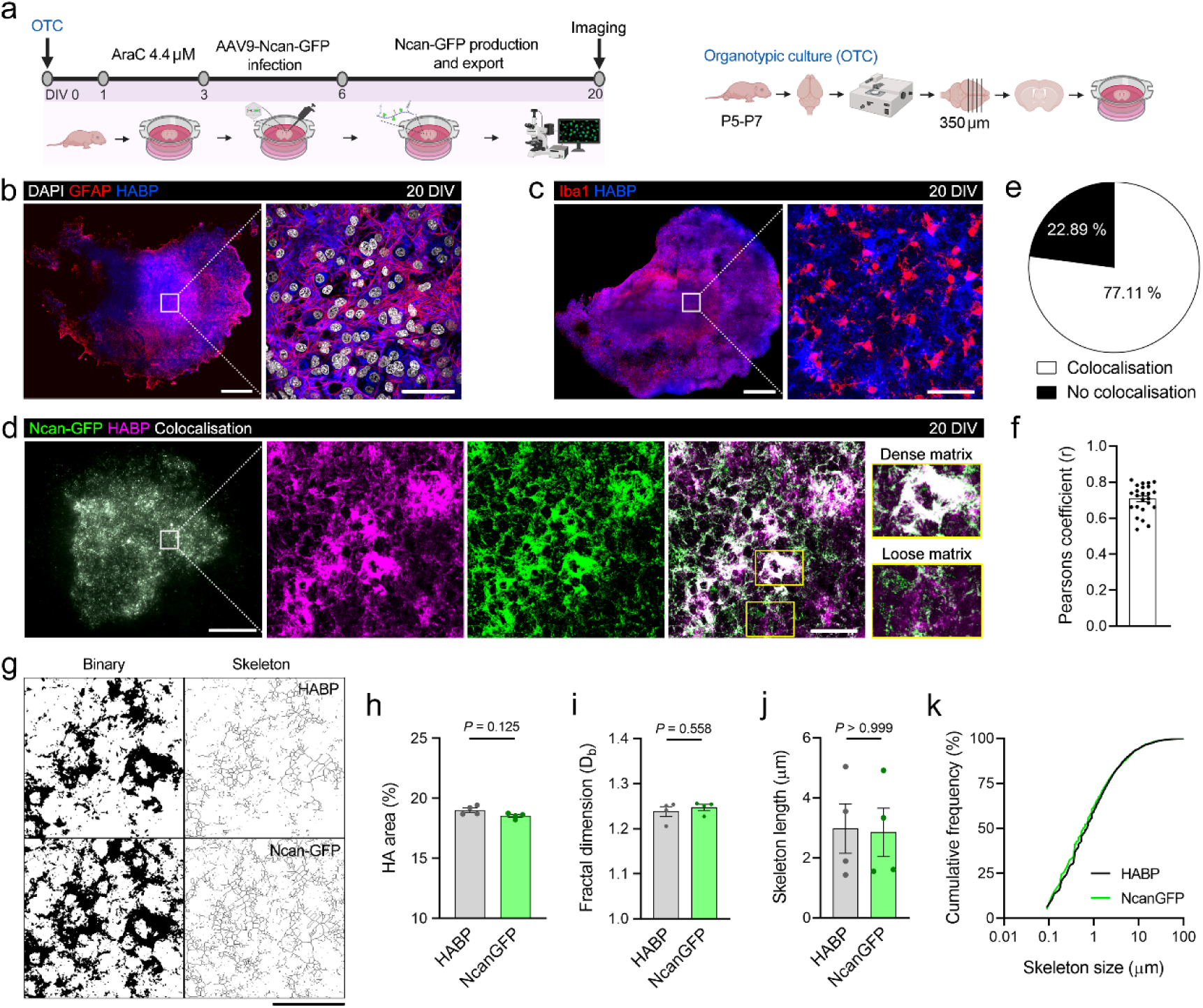
AAV-Ncan-GFP labels HA and reveals iECM heterogeneity in OTC slices. **a)** Experimental timeline of AAV-Ncan-GFP infection and imaging in OTC slices. **b, c)** Confocal tile-scan of DIV 20 OTC slices showing distribution of HA (HABP) in the context of astrocytes (GFAP) and microglia (Iba1), responsible for ECM homeostasis in brain tissue. Scale bars, 1 mm and 50 µm (insets). **d)** Confocal tile-scan of DIV 20 OTC slice showing distribution of Ncan-GFP vital labeling of HA, in colocalization with HA labeling (HABP) in fixed OTC tissue. Insets show colocalization in areas of dense ECM, with lower colocalization in regions with loose ECM. Scale bar, 1 mm and 50 µm (inset). **e, f)** Over 70 % (**e**) of HA Ncan-GFP labeling colocalizes with HABP staining (Pearson’s coefficient, r = 0.707). **g)** Binary and skeletonized images of the HA network used to quantify HA area and structure. Note that Ncan-GFP and HABP patterns are similar. Scale bar, 50 µm. **h)** No significant difference was found in HA levels when labeled with Ncan-GFP or HABP (Wilcoxon matched-pairs test, *n* = 4 OTC, 8 crops each). **I)** Fractal dimension analysis of the HA network labeled by Ncan-GFP shows similar matrix interconnectivity than HABP staining (two-tailed Student’s paired t-test, *n* = 4 OTC, 8 crops each). **j, k)** Mean (**j**) and cumulative distribution (**k**) of HA skeleton lengths indicate similar cable-like structures of the brain ECM with both Ncan-GFP and HABP labeling (Kolmogorov-Smirnov test, *n* = 4 OTC, 8 crops each).

We first characterized the HA network in fixed OTC slices by immunofluorescence. At DIV 20, HABP staining revealed the typical profuse and cable-like interconnected distribution of brain interstitial HA reported *in vivo* ^15,29^. We had previously observed that glial cells are critical for HA homeostasis ^15,30^, therefore we performed HABP staining in conjunction with glial cell markers. HA was stained alongside GFAP-positive astrocytes (**Figure 2b**) and Iba1-positive microglia (**Figure 2c**), both of which are present in significant numbers in the OTC tissue. From DIV 15 onward, AAV-Ncan-GFP vital labeling revealed an HA distribution comparable to that observed with HABP, the standard labeling method for HA. HA cable-like structures, often described in the literature ^31^, became prominent at DIV 20 when AAV expression was maximal, allowing the distinction between areas of dense and loose iECM (**Figure 2d**). Areas of dense HA showed a greater colocalization than loose iECM regions. Quantification of HA area revealed that more than 70% of Ncan-GFP labeling colocalizes with HABP staining (Pearson’s coefficient r = 0.707; **Figure 2e**, **f**), with no significant differences detected in HA content between both labeling methods (18.55 ± 0.18% Ncan-GFP; 19.03 ± 0.12% HABP; **Figure 2g**, **h**).

Although HA is a linear polysaccharide, it forms an interconnected network through cross-linking by hyalectans and tenascins, resulting in supramolecular complexes of varying stability ^32^. Measuring HA cross-linking provide, thus, an estimation of matrix interconnectivity and organization ^15,30^. Therefore, we quantified the HA network complexity labeled with Ncan-GFP using fractal analysis (Db) ^33^, and found similar values to those obtained with HABP staining (1.25 ± 0.01% Ncan-GFP, 1.24 ± 0.007% HABP; **Figure 2i**), indicating that the HA network complexity reported by our vital labeling is equivalent to that observed with conventional HA labeling methods. Furthermore, skeleton analysis of Ncan-GFP labeling revealed HA cables of a similar length to those labeled with HABP (median = 0.58 µm Ncan-GFP, 0.7 µm HABP; **Figure 2j**, **k**). These results validate Ncan-GFP labeling of HA, positioning it on par with current HA labelling methods, with the added advantage of being genetically encoded.

To further validate the utility of the probe, we explored Ncan-GFP labeling at submicron resolution, shifting our focus to a finer-scale brain structure: neuronal spines. At these scales, fixation artifacts become more pronounced, potentially altering the delicate structure of the iECM. To avoid such artifacts, we performed time-lapse confocal imaging with a water-immersion objective, allowing us to visualize the fine structure of iECM and dendritic spines in live OTC slices. We co-infected OTC slices with AAV-Ncan-GFP and AAV-SYN-tdTomato, enabling real-time visualization of both HA and neurons (**Figure 3a**). The probe effectively captures the spatial heterogeneity of the iECM at micrometer resolution with exceptional photostability. This allowed for precise visualization of iECM in relation to neuronal structures, revealing that membrane-bound HA is heterogeneously distributed along the neuronal soma (**Figure 3b**, **c**), dendrites (**Figure 3d**, **e**) and spines (**Figure 3f**, **g**).

**Fig. 3.**
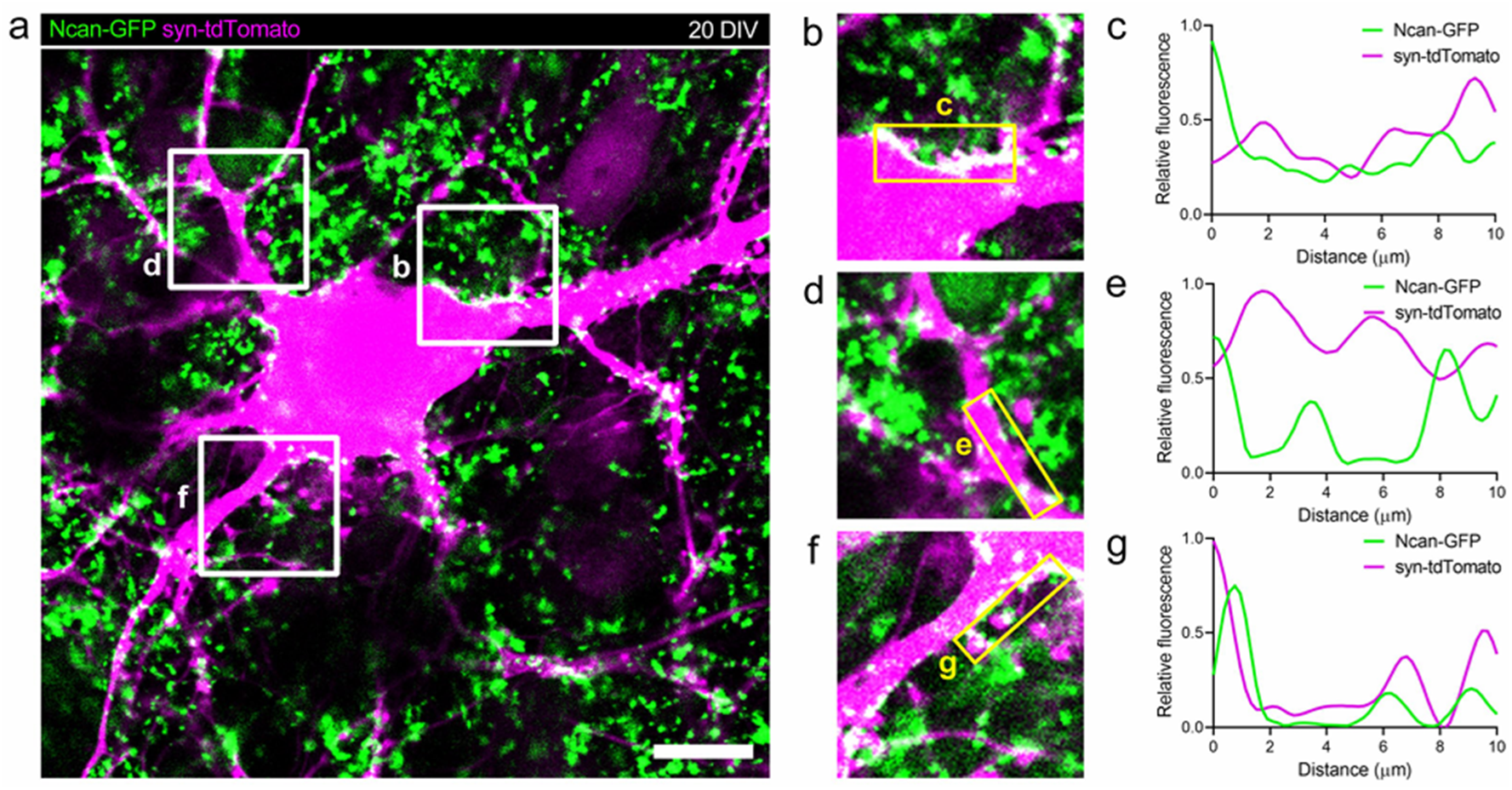
AAV-Ncan-GFP reveals iECM heterogeneity at submicron resolution. **a)** High-resolution live confocal image showing distribution of HA (Ncan-GFP) surrounding a cortical neuron (Syn-tdTomato) in OTC slices at DIV 20. Scale bar, 20 µm. **b, d, f)** Insets from (**a**) showing ECM distribution around neuronal soma (**b**), dendrites (**d**) and dendritic spines (**f**). **c, e, g)** Fluorescence intensity profiles from (b, d, f) indicate that membrane-bound HA is heterogeneously distributed along the neuronal soma (**b**) and dendrites (**d**), and spines (**f**).

### AAV-Ncan-GFP labels extracellular hyaluronan in live tissue

Since the Ncan-GFP probe is produced intracellularly, but its target (HA) is located in the ECS, we next verified that Ncan-GFP vital labeling is mostly extracellular. To visualize the ECS, we employed confocal-shadow imaging (COSHI) ^24^, which provides detailed optical access in real-time to the anatomical complexity and dynamics of living brain tissue at the micrometric scale using a relatively simple setup. We then imaged, in a confocal microscope, live OTC slices with both AAV-Ncan-GFP-mediated HA labeling and AlexaFluor-594 labeling of the interstitial fluid, to visualize the brain ECM and ECS simultaneously (**Figure 4a**). To test if the ECS and ECM were discernible at confocal diffraction-limited resolution, we labeled the ECM using the “pan-ECM” method, a recently developed chemical labeling technique that binds covalently to both extracellular and membrane-bound proteins, instead of glycans ^34^. Pan-ECM labeling was found all along the ECS compartment, visualized in dark at the inverted image, with sufficient resolution to discern extracellular from intracellular structures (**Figure 4b**). Ncan-GFP vital staining displayed a similar extracellular distribution, although it occupied a fraction of the total ECS width (**Figure 4c**, **d**). This difference might stem from AAV-Ncan-GFP labeling only interstitial HA (iECM), while pan-ECM targets proteins both at the extracellular and membrane compartments, providing a staining similar to the ECS labeling performed in shadow imaging ^34^.

**Fig. 4.**
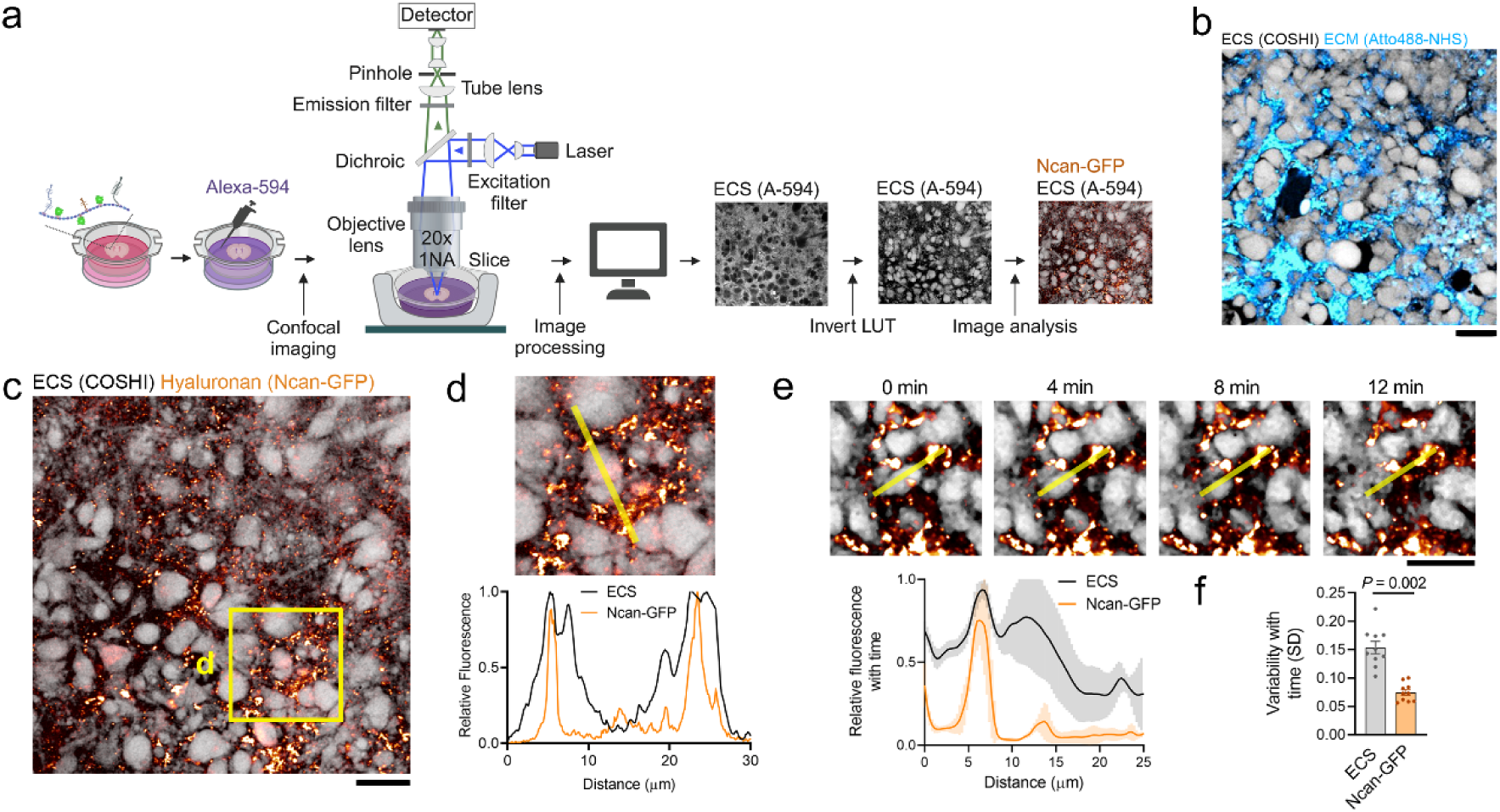
AAV-Ncan-GFP labels extracellular HA in live tissue. **a)** Schematic diagram of confocal shadow imaging (COSHI) alongside Ncan-GFP labeling of the brain ECM. AlexaFluor-594 dye is diluted in culture medium allowing to diffuse through the extracellular compartment for 1h. OTC slices are then imaged at 37°C in an incubation chamber with 40x water immersion objective. **b)** COSHI image of OTC tissue showing ECM protein distribution (Atto488-NHS) with the “pan-ECM” staining method, alongside the ECS (in dark). Scale bar, 20 µm. **c)** Confocal shadow image (AlexaFluor-594) of OTC slice tissue showing brain iECM distribution (Ncan-GFP) alongside ECS (in dark). Scale bar, 20 µm. **d)** Inset from (**c**). The fluorescence intensity profile confirms that Ncan-GFP labeling is localized in the ECS. **e)** Real-time series of COSHI + Ncan-GFP imaging in a dynamic area in live OTC tissue. The fluorescence intensity profile shows the ECS (in dark) dynamically changing with time, while brain ECM (Ncan-GFP) remains mostly stable at short timeframes. Scale bar, 20 µm. **f)** Variability with time is significantly higher in the ECS compartment, compared to brain ECM (Ncan-GFP), quantified as the mean standard deviation of fluorescence intensity profile with time (Wilcoxon test, *n* = 3 OTC, 3 crops each).

A key advantage of Ncan-GFP is that it binds hyaluronan non-covalently, which does not alter the cross-linking of the ECM, allowing for the observation of its dynamics in a more natural state. Time-lapse recording of both COSHI and AAV-Ncan-GFP staining revealed, however, that while the ECS changes dynamically within a few minutes, the brain iECM remains mostly stable over this short period, as indicated by the mean standard deviation of all time frames (0.15 ± 0.01% ECS vs 0.07 ± 0.005% Ncan-GFP; **Figure 4e**, **f**). Taken together, these results confirm the efficacy of AAV-Ncan-GFP as a tool for labeling the extracellular HA network in real-time in live brain tissue.

### AAV-Ncan-GFP labeling is sensitive to ECM impairment

Next, we went further in the validation of the probe by testing its sensitivity to hyaluronidase (Hyase). Since the bond between Ncan-GFP and HA is non-covalent, we hypothesized that HA degradation would dismantle Ncan anchor points at HA, leading to the dilution of the probe into the interstitial fluid and a decrease in fluorescence. We followed Hyase-mediated HA matrix disruption in real-time by confocal microscopy in OTC slices, and detected a drastic reduction in Ncan-GFP labeling 15 min after Hyase treatment (**Figure 5a**, **b**). Since Ncan-GFP is produced intracellularly and Hyase acts only in the ECS, we interpreted the remaining high intensity values after Hyase-mediated degradation as corresponding to intracellular Ncan-GFP. This labeling not only displayed a cell-like morphology but also showed intensity patterns consistent with a higher degree of compartmentalization, typical of the intracellular compartment (**Figure 5c**). Finally, both Ncan-GFP and HABP areas were significantly reduced after HA impairment, first by Hyase-mediated HA degradation, but also when OTC slices were treated with 4-methylumbelliferyl glucuronide (4-MUG), an inhibitor of HA synthesis used in *in vitro* paradigms ^35^ (**Figure 5d**). These results further confirm the specificity of AAV-Ncan-GFP to label extracellular HA, while also highlighting its potential for monitoring ECM impairment.

**Fig. 5.**
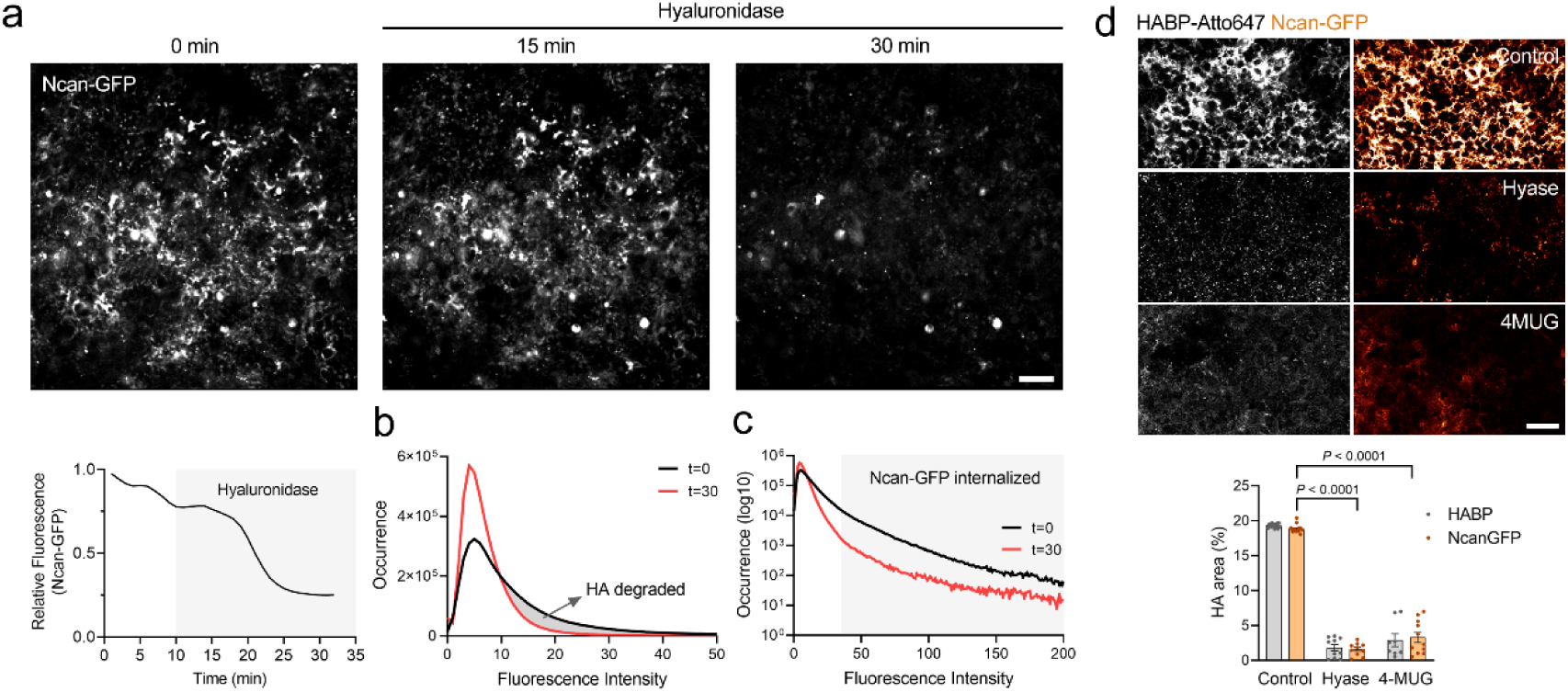
AAV-Ncan-GFP labeling is sensitive to ECM impairment. **a)** Real-time series and relative fluorescence histogram of confocal images of the HA network (Ncan-GFP), show brain ECM disruption 15 min after hyaluronidase treatment in OTC slices. Scale bar, 20 µm. **b)** Fluorescence intensity histogram of Ncan-GFP labeling before (t = 0 min) and after (t = 30 min) Hyase treatment shows HA breakdown. The empty area between the curves corresponds to the HA fraction degraded. **c)** Logarithmic fluorescence intensity histogram shows that the higher intensity values observed after Hyase treatment (t = 30 min) correspond to intracellular Ncan-GFP, as these values are also present before HA disruption (t = 0 min). **d)** Confocal micrographs and quantification of HABP and Ncan-GFP staining in OTC slices after Hyase-mediated HA degradation and 4-MUG-mediated inhibition of HA synthesis demonstrate that Ncan-GFP effectively reports a decrease in HA content (Two-way ANOVA with Tukey’s post-hoc test, *n* = 3 OTC, 3 crops each). Scale bar, 20 µm.

### AAV-Ncan-GFP labels both iECM and PNNs in vivo

Having validated AAV-Ncan-GFP as a real-time probe for labeling HA in OTC slices, we next assessed its effectivity *in vivo*. We injected AAV-Ncan-GFP into the motor cortex (M1) of adult mice and performed immunofluorescence and confocal imaging 30 days post-injection (dpi), when Ncan-GFP expression peaks (**Figure 6a**). We observed extensive Ncan-GFP labeling of HA in the brain parenchyma, with high colocalization with HABP and a similar staining pattern, revealing the local heterogeneity of HA (**Figure 6b**, **c**). Interestingly, Ncan-GFP labeled not only HA in the iECM, but also in PNNs, which are specialized ECM structures surrounding mainly parvalbumin-positive interneurons ^10^ and containing a dense and highly organized HA. We confirmed Ncan-GFP staining of PNNs by co-staining with the CSPG-specific lectin WFA. We observed precise colocalization of Ncan-GFP and WFA staining (**Figure 6d**, **e**). These findings demonstrate that AAV-Ncan-GFP labels *in vivo* both the brain iECM, but also PNNs, enabling extensive survey of the local brain ECM.

**Fig. 6.**
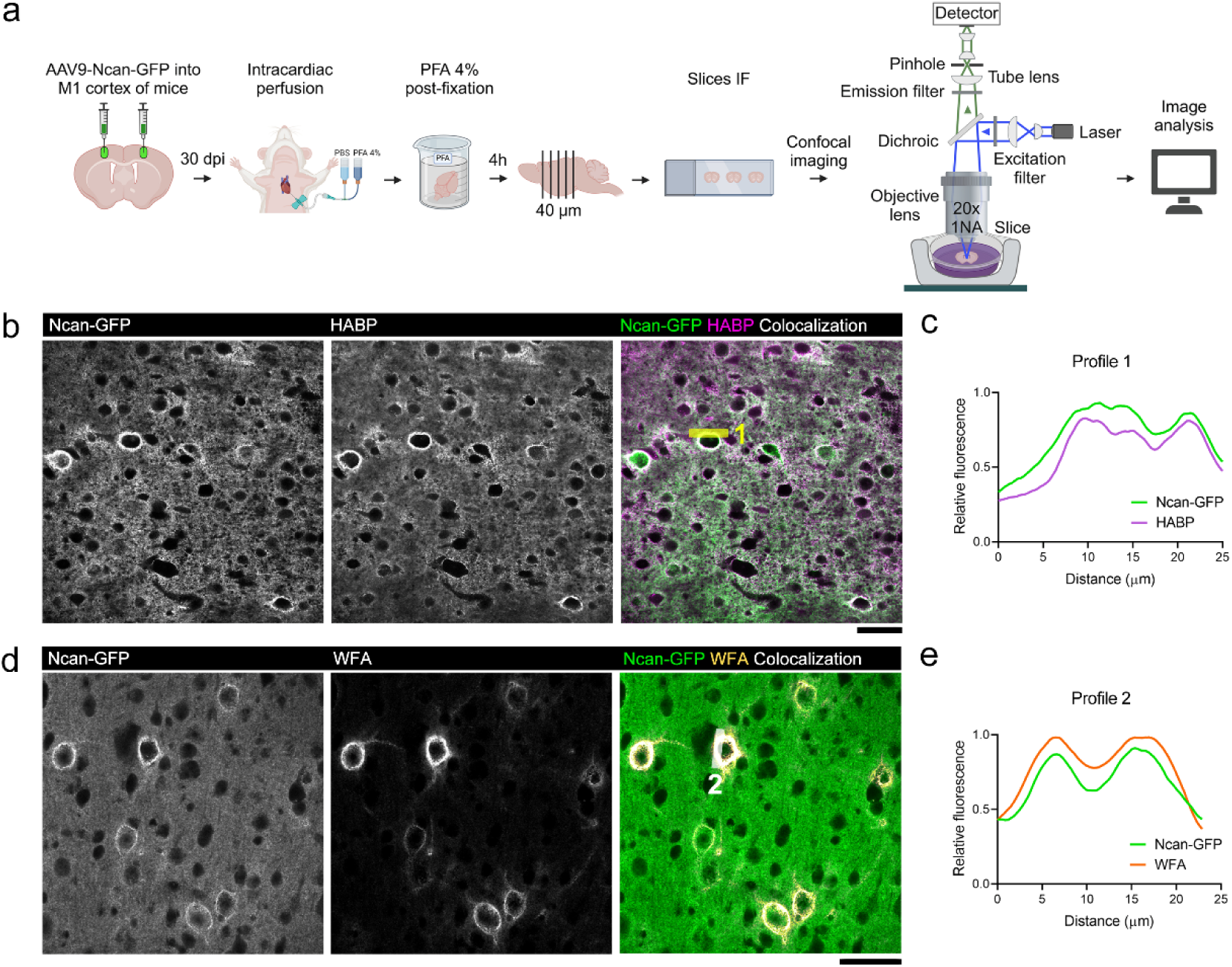
AAV-Ncan-GFP labels iECM and PNNs *in vivo*. **a)** Schematic diagram of AAV-Ncan-GFP inoculation, fixation and imaging of tissue sections of adult mice in confocal imaging. **b)** Confocal micrographs from the motor cortex (M1) of AAV-Ncan-GFP-injected adult mice at 30 dpi, showing a similar staining pattern of iECM with AAV-Ncan-GFP and standard HA staining with HABP. Scale bar, 50 µm. **c)** Fluorescence intensity profile from (**b**) showing colocalization between the two staining methods. **d)** Confocal micrographs of AAV-Ncan-GFP-injected adult mouse cortex, showing Ncan-GFP labeling of PNNs, colocalizing with PNN’s specific staining WFA in fixed tissue. Scale bar, 50 µm. **e)** Fluorescence intensity profile 2 from (**d**).

### AAV-Ncan-GFP enables visualization of brain ECM dynamics in vivo

As a final validation of our tool, we tested AAV-Ncan-GFP for live imaging of the brain ECM in vivo in anesthetized animals. Since Ncan-GFP expression peaks at 30 dpi, we injected the vector into the somatosensory cortex of adult mice and simultaneously implanted a cranial window, to be able to image 30 days after the surgery with minimal inflammation. We then performed two-photon microscopy to interrogate the brain ECM in real-time in the mouse cortex (**Figure 7a**). The cranial window approach allows deep imaging of the live brain at different timeframes, ranging from minutes to days ^36^. AAV-Ncan-GFP revealed the spatial distribution of brain iECM in layers II-III of mouse cortex at 27 dpi (**Figure 7b**), with an extensive extracellular labeling despite only a few sparse neurons and glia being transfected, which also exhibited intracellular fluorescence. To assess iECM dynamics, we then performed timelapse imaging of the HA network (Ncan-GFP) and observed that HA local content remained stable in the *short-term*, with minimal variation in its fluorescence profile after 18 minutes (**Figure 7c**, **d**). To assess iECM dynamics in the *long-term*, we imaged the same brain region through the cranial window, following iECM dynamics during 15 days, until 42 dpi (**Figure 7e**), although the expression of the probe was stable even until 60 dpi (data not shown). Time-lapse imaging of Ncan-GFP revealed that the homeostatic HA matrix exhibits greater variability in the long term (days) than in the short-term (minutes), while still maintaining a stable local distribution (**Figure 7f**).

**Fig. 7.**
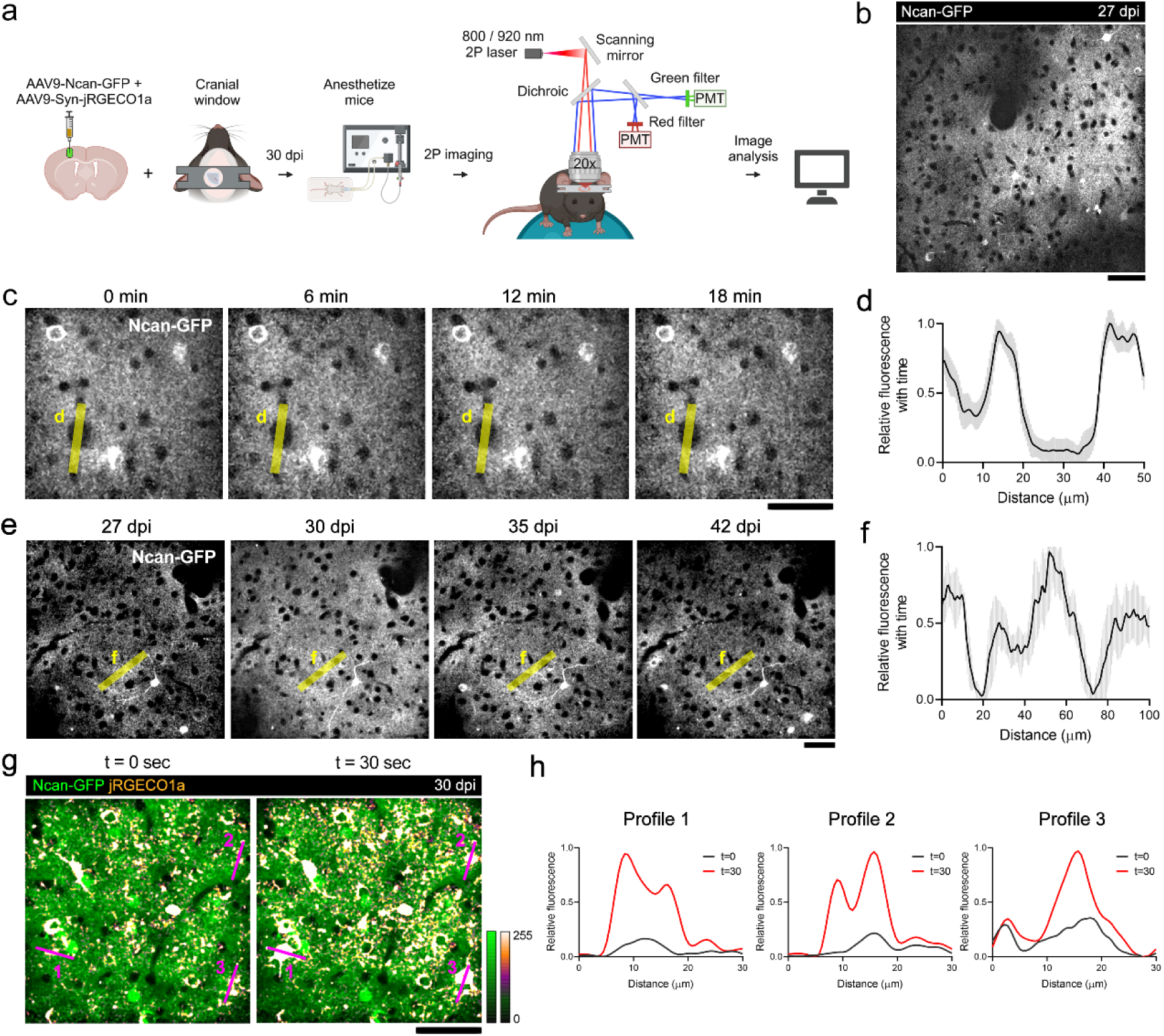
AAV-Ncan-GFP *in vivo* enables detection of brain ECM dynamics. **a)** Schematic diagram of two-photon *in vivo* imaging of AAV-Ncan-GFP through a cranial window in anesthetized mice. **b)** Two-photon image of brain iECM (Ncan-GFP) at 200 μm depth (layer II-III) in somatosensory cortex of adult mice 27 dpi. Scale bar, 50 µm. **c)** Two-photon time-lapse imaging of adult mouse cortex with Ncan-GFP brain ECM labeling. Images were acquired for consecutive minutes with minimal photobleaching. Labeling of both iECM and PNN can be observed. Scale bar, 50 µm. **d)** Fluorescence intensity profile from (**c**) shows brain iECM and PNNs (Ncan-GFP) remain mostly stable in the short-term (18 min). Low standard deviation (in gray) indicates HA network does not change within this timeframe. **e)** Longitudinal imaging of somatosensory cortex (layers II-III) with Ncan-GFP labeling from 27 to 42 dpi. Images was performed in consecutive weeks in adult mice. Scale bar, 50 µm. **f)** Fluorescence intensity profile from (**e**) shows brain ECM (Ncan-GFP) remain mostly stable in the long-term (along 15 days) in homeostatic conditions, with few variations in Ncan-GFP fluorescence intensity. Higher standard deviation (in gray) was observed when compared with short-term imaging (**d**). **g)** Two-photon imaging of neuronal Ca^2+^ activity (AAV-Syn-NES-jRGECO1a) in the context of brain ECM (Ncan-GFP) in the cortex. Images were recorded during 10 minutes (1 Hz) in adult mice at 0.5% anesthesia. Scale bar, 50 µm. Color bars represent fluorescence intensity. **h)** Fluorescence intensity profiles from (**g**) show intracellular calcium fluctuations in neurons.

Next, we explored whether AAV-Ncan-GFP is compatible with other *in vivo* probes, such as those used to study cell dynamics. To test this, we used AAV-Syn-NES-jRGECO1a, a genetically encoded calcium indicator for measuring activity in neuronal populations ^37^. We co-injected this vector with AAV-Ncan-GFP in the somatosensory cortex of adult mice and implanted a cranial window on top for two-photon time-lapse imaging *in vivo*. At 30 dpi, spontaneous neuronal Ca^2+^ activity (jRGECO1a) was recorded during 10 minutes (1 Hz) in the context of the HA matrix in vivo (**Figure 7g**). Fluorescence intensity profiles revealed neuronal increase in fluorescence due to spontaneous Ca^2+^ transients (**Figure 7h**). Taken together, these results demonstrate that AAV-Ncan-GFP can be used to study brain ECM dynamics in living mice across both short and long timeframes, alongside cellular dynamics such as dendritic spines (Figure 3) or calcium activity (Figure 7). This vector thus enables tracking neuronal (and plausibly glial) dynamics within the context of brain iECM or PNNs in vivo.

## DISCUSSION

Designing a tool that labels the brain ECM structure in real-time represents a biological and technological challenge, as its main component, HA, is a glycan polymer that is neither immunogenic nor genetically encoded. Despite some ECM staining methods have been reported ^18,34,38^, no tools are currently available for live imaging of the HA matrix structure and dynamics in mice, the most commonly used species in neuroscience research. Here we demonstrated stable expression of AAV-Ncan-GFP probe in organotypic brain slices, as well as *in vivo* in living mice, labeling HA in both PNNs and iECM. We also validated HA labeling through colocalization with HABP and WFA, and sensitivity to hyaluronidase and HA synthesis inhibition, further confirming the probe extracellular localization by confocal shadow imaging. We were able also to conduct, for the first time, longitudinal time-lapse imaging of ECM dynamics in live mice. Furthermore, we combined AAV-Ncan-GFP with calcium transient imaging in vivo, providing a real-time local map of the iECM alongside neural function.

The brain ECM has traditionally been visualized in still images using immunofluorescence with HA-binding proteins ^39^. But fixation not only prevents the observation of dynamic changes but can also alter the pattern of HA staining ^40^. Here, we validate a new tool that provides a structural view of the HA matrix in live tissue, allowing visualization of the brain ECM in its native form. AAV-Ncan-GFP staining reveals the spatial heterogeneity of HA at both micron and submicron resolution, enabling the exploration of the ECM alongside microscopic structures such as glial processes or neuronal dendrites. Furthermore, since the probe is based on a fluorescent protein, it is compatible with super-resolution techniques such as STED ^41^, allowing the investigation of ECM alongside nanoscale events at the synapse, including exocytosis ^42^ and receptor mobility ^43^. Since the iECM is a primary determinant of ECS structure and diffusional properties ^1,4^, AAV-Ncan-GFP could be combined with ECS exploration techniques such as super-resolution shadow imaging (SUSHI) ^23,24^ and carbon nanotube ^44,45^ or nanoparticle ^46^ tracking to further investigate the interplay between the ECS and its ECM scaffold in live tissue. We have successfully tested AAV-Ncan-GFP in combination with neuronal fine structures and calcium transients, both in OTC slices and *in vivo*, demonstrating its utility for exploring brain spatiotemporal dynamics within the context of the ECM. The probe, thus, opens new opportunities to improve our understanding of ECM’s role in brain dynamics.

Regarding its specificity, AAV-Ncan-GFP renders an HA distribution comparable to that observed with HABP, the current gold standard for labeling the iECM. However, AAV-Ncan-GFP exhibits over 70% colocalization with HABP, which, while high, is not a perfect match. This discrepancy is likely due to incomplete viral transduction across the tissue, as only well-transfected areas will produce sufficient amounts of the fluorescent hyalectan to properly label the HA network. It is also plausible that exogenous HABP staining is not fully labelling the parenchyma, potentially missing smaller ECS microdomains, and therefore not binding to the same HA areas as the endogenous (secreted) Ncan-GFP probe. Despite this limitations, AAV-Ncan-GFP effectively reveals areas of both loose (iECM) and condensed (PNNs) HA, highlighting its value in reporting local HA heterogeneity. It should also be noted that although the probe is designed with an export signal for release into the ECS, a small portion is still intracellular at any given time. The higher fluorescence intensity values correspond to this intracellular hyalectan, as it persists even after HA matrix disruption with Hyase (Fig 4). This bright intracellular GFP is easy to differentiate, even at low resolution, from the diffuse extracellular Ncan-GFP. This is because the GFP inside cells is not only more concentrated than in the ECS, but partially loses fluorescence yield in the extracellular milieu ^47^. Researchers using the probe should be aware of these limitations and take measures to overcome them, such as selecting areas with sufficient vector transduction and excluding sparse intracellular GFP based on morphology or fluorescence intensity values.

On the same note, AAV-Ncan-GFP imaging should be conducted when expression is maximal, to ensure proper labeling of the ECM. We observed that in cortico-striatal OTC slices, Ncan-GFP peaks at DIV 15, and we were able to image the ECM in OTCs up to DIV 30. In contrast, *in vivo*, widespread labeling of the ECM is achieved after 30 days, and it persists until 60 dpi, where it begins to decline. The long-term stability of the staining allowed us to confirm that, in fact, the iECM remains mostly stable over time, with greater fluctuations observed *in vivo* compared to OTC slices. This could be due to increased production of fluorescent hyalectan *in vivo*, or possibly because the HA meshwork becomes denser with age ^48,49^. This ability of Ncan-GFP to report HA accumulation, combined with its sensitivity to ECM disruption (Figure 4) would enable researchers to explore the dramatic changes that the ECM undergoes in pathological conditions. For instance, it can be used to follow HA degradation in neurodegeneration ^15^, or observe in real-time accumulated HA at the glial scar ^30^.

Outside of the brain, live visualization of ECM has primarily been conducted with second harmonic generation (SHG) under two-photon excitation. However, this method predominantly illuminates fibrillar collagen ^50^, which is largely absent in brain ECM. A newer method, termed “pan-ECM”, utilizes protein-reactive dyes such as Atto488-NHS, which covalently tag extracellular amines ^34^. While this approach labels the ECM by tagging non-diffusible extracellular proteins, it also stains all proteins exposed to the extracellular environment, including membrane-bound receptors or ion channels. Furthermore, it remains unclear whether the covalent nature of the bond in “pan-ECM” labeling further cross-links the matrix meshwork, potentially limiting the study of ECM dynamics. Still, it is a useful method for acutely tagging the protein fraction of the ECM. In contrast, Ncan-GFP labels the glycan portion and, due to its genetically-encoded nature and non-covalent binding, enables chronic ECM staining and exploration of ECM dynamics, including ECM disruption. Thus, both methods are complementary and, when combined, may provide a more comprehensive visualization of the extracellular scaffold.

AAV-Ncan-GFP, in its current form, utilizes the ubiquitous promoter CMV to drive Ncan-GFP expression, ensuring synthesis and secretion by many cell types for widespread ECM labeling. Future iterations of the probe could incorporate cell-specific promoters or enhancers, such as those targeting PV-positive interneurons, for instance, for PNN-specific labelling, or the GFAP promoter to focus on the iECM within the glial scar. Additional developments might include switching to red fluorophores, enabling compatibility with green reporters like GCaMP or GFP-based cellular markers. Integrating a cysteine-free GFP variant ^51,52^ could further enhance probe brightness in the oxidative extracellular environment. We anticipate that the probe will continue evolving over time to address diverse biological questions and experimental paradigms.

Over the past decades, the development of fluorescent-labelling techniques to tag brain cells, ranging from diffusible dyes to genetically-encoded fusion proteins, has revolutionized our understanding of brain function, enabling the visualization of dynamic processes such as dendritic spine plasticity and microglial process motility. ECM fluorescent reporters such as AAV-Ncan-GFP will add a missing piece of the puzzle, the labelling of the iECM *in vivo*. Most of our understanding of the ECM’s influence on brain function stems from studies on PNNs ^2,53^, an important yet limited perspective of the broader ECM. Imaging the iECM, in combination with cell-specific reporters and ECM-modifying paradigms, will provide new insights on how, and especially *where*, this ubiquitous structure, which envelops nearly every cell membrane, influences brain dynamics. This includes not only its effects on brain cells but also the ECS itself, an interplay that remains largely unexplored. This tool will thus provide an unprecedented opportunity to make a leap forward in our understanding of extracellular control of cellular function, beyond the realm of PNNs and inhibitory circuits.

## ACKNOWLEDGEMENTS

The authors would like to thank Valeria Montoya and Laura Escobar for their invaluable technical support. We thank Dr. Kelly Smith (University of Queensland, Australia) for providing the original Ncan-GFP plasmid for zebrafish. We thank Dr. Alejandro Carretero (Achucarro Basque Center for Neuroscience) for the AAV-hSYN-tdTomato plasmid. F.N.S. acknowledges funding from the Spanish Ministry of Science and Innovation (PID2020-115896RJ-I00, PID2023-152586OB-I00 and RYC2021-032602-I co-funded by NextGeneration EU/PRTR). M.F.B. was supported by a PhD fellowship from the Basque Government.

## AUTHOR CONTRIBUTIONS

M.F.B. conducted experiments, collected and analyzed data and prepared the figures. M.A. assisted in AAV-inoculation, cranial window implantation and *in vivo* two-photon imaging. N.D. designed the AAV-Ncan-GFP vector. N.D. and L.A.L. produced the AAV-Ncan-GFP viral particles. A.M. provided infrastructural support and scientific feedback. F.N.S. conceived and supervised the study and secured funding. M.F.B. and F.N.S. wrote the paper.

## METHODS

### Animals

Male and female wild-type C57BL/6J mice (Janvier) were used for *in vivo* experiments. Sprague Dawley rat pups from P5 to P7 were used for organotypic brain slices. Animal handling and experimental procedures were performed in accordance with the European Communities Council Directive and approved by the Ethics Committee of the University of the Basque Country (UPV/EHU) under licenses M20/2022/243 and M20/2023/030. All possible efforts were made to minimize animal suffering and the number of animals used.

### Plasmid construction and AAV vector production

The expression plasmid pAAV-CMVie-ss-Ncan-GFP-WPRE was generated by deletion of the synapsin promoter and synuclein encoding region from plasmid pAAV-SynP-synA53T-WPRE ^54^ (provided by Dr. V. Baekelandt) and insertion of the CMVie94 promoter and ssNcan-EGFP sequences from plasmid pCS2-HA-GFP ^18^ (kind gift from Dr. Kelly Smith). AAV vectors were produced and purified as previously described ^55^. In brief, AAV vectors were produced by polyethyleneimine (PEI)-mediated triple transfection of low passage HEK-293T /17 cells (ATCC; cat number CRL-11268). The AAV expression plasmids were co-transfected with the adeno helper pAd Delta F6 plasmid (Penn Vector Core, cat # PL-F-PVADF6) and AAV Rep Cap pAAV2/9 plasmid (Penn Vector Core, cat # PL-T-PV008). Cells were harvested 72h post transfection, resuspended in lysis buffer (150 mM NaCl, 50 mM Tris-HCl pH 8.5) and lysed by 3 freeze-thaw cycles (37°C/-80°C). The cell lysate was treated with 150 U/ml Benzonase (Sigma) for 1 hour at 37°C and the crude lysate was clarified by centrifugation. Vectors were purified by iodixanol step gradient centrifugation, and concentrated and buffer-exchanged into Lactated Ringer’s solution (Baxter, Deerfield, IL) using vivaspin20 100 kDa cut off concentrator (Sartorius Stedim). Titrations were performed at the Transcriptomics Platform (Neurocentre Magendie, INSERM U862, Bordeaux, France). The genome-containing particle (gcp) titer was determined by quantitative real-time PCR using the Light Cycler 480 sybr green master mix (Roche, cat # 04887352001) with primers specific for the AAV2 ITRs (fwd 5′-GGAACCCCTAGTGATGGAGTT-3′; rev 5′-CGGCCTCAGTGAGCGA-3′) ^56^ on a Light Cycler 480 instrument. Purity assessment of vector stocks was estimated by loading 10 µl of vector stock on 10% SDS acrylamide gels, total proteins were visualized using the Krypton Infrared Protein Stain according to the manufacturer’s instructions (Life Technologies).

### Organotypic brain slices

Organotypic cultures were prepared according to the interface culture method ^25^, with minor modifications ^57^. Briefly, cortico-striatal slices were obtained from postnatal day 5-7-old Sprague-Dawley rat pups. Animals were quickly decapitated, and the brains sliced on a McIlwain tissue chopper to generate coronal slices of 350 µm thickness. Brain slices were separated in cold sterile HBSS without calcium and magnesium (#14175095, Thermo Fisher). Within 20 min maximum, slices were transferred onto sterilized Millipore 0.4 µm culture inserts (PICM0RG50; Merck Millipore) held in a 6-well plate with preheated culture medium (50% Minimum Essential Medium Eagle’s (MEM), 25% Earle’s Balanced Salt Solution (EBSS), 25% Horse Serum, 36 mM glucose, B27 Supplement, GlutaMAX, 50 U/ml Penicillin-Streptomycin and 1.25 µg/ml Fungizone (Amphotericin B); all from GIBCO). The following day, antimitotic Cytosine β-D-arabinofuranoside (AraC) 4.4 µM was added to the feeding medium. At DIV 3, AraC was removed and slices were infected with AAV-Ncan-GFP 10^12^ vg/ml or co-infected with AAV9-Syn tdTomato 10^10^ vg/ml over-weekend. 1 µl of viral particles were added over the OTC slice, and from this point, culture medium was replaced every two days. Organotypic slices were cultured up to 20 days at 35°C / 5%CO_2_ and imaged in the same culture medium from DIV 15 onwards. A detailed protocol of the OTC culture and AAV-Ncan-GFP inoculation has been published in *protocols.io* ^28^.

### Live imaging in OTC slices

Cortico-striatal OTC slices infected with AAV-Ncan-GFP were imaged starting at DIV 15 in a Leica Stellaris 5 microscope equipped with a heated chamber (OkoLab), a vertical 40X water-immersion objective (NA = 0.8). Imaging was performed at 37°C in the same OTC medium. Images (pixel size = 100 nm) were performed at 1 min intervals in the same z plane, controlling the drift in z every 5 min. We used minimum power and maximum gain to minimize phototoxicity and photobleaching. For “pan-ECM” staining, 0.1 mM of Atto-488-NHS was added to the slices 1h prior to imaging. For COSHI, the ECS was previously stained with 40 μM AlexaFluor 594 (A33082, ThermoFisher), as described elsewhere ^24^. For visualization of cellular structures in COSHI, the inverted image was used.

### Stereotaxic surgery

3–5-month-old C57BL/6J mice of both sexes were bilaterally injected into the motor cortex (M1) with 500 nl of AAV-Ncan-GFP 10^13^ vg/ml, by stereotactic surgery (coordinates from bregma: +1 mm anteroposterior, +/-1.5 mm mediolateral, and -1.5 mm dorsoventral) under deep isoflurane anesthesia. Injection was performed with a 30-gauge Hamilton syringe coupled to a microinjection pump at a rate of 60 nl/min. The needle was left in place for 10 min to avoid AAV leakage. Animals were injected IP with buprenorphine (0.1 mg/kg) during the surgery and every 12 h (twice) to minimize pain.

### Immunofluorescence

Mice were anesthetized with pentobarbital (100 mg/kg ip) and intracardially perfused with 4% paraformaldehyde (PFA) in 0.1M PB. Brains were post-fixed for 3 h in 4% PFA and 40 μm-thick vibratome coronal sections were collected. For visualization of the HA matrix or CSPGs, slices were blocked with 1% bovine serum albumin (BSA) in PBS with 0.1% saponin for 1 h, followed by streptavidin-biotin blocking kit (Vector Labs) for 20 min, and incubated over-weekend with either biotinylated-hyaluronic acid binding protein (HABP from bovine nasal cartilage, Merck-Millipore) or biotinylated Wisteria Floribunda Agglutinin (WFA, Vector Labs) diluted in blocking solution. Staining was revealed with Streptavidin-Atto647N (Sigma-Aldrich), with minimum non-specific labeling due to appropriate blocking of endogenous biotins ^15^.

Cortico-striatal organotypic slices were fixed at DIV 20 for 1 hour with 4% PFA in 0.1M PB. Immunofluorescence was performed similarly to mouse tissue sections. Tissue sections and OTC slices were mounted on #1.5 coverslips with Mowiol + DABCO and left to dry overnight in darkness. Confocal images were obtained in a Leica Stellaris microscope, maintaining image acquisition settings (laser power, AOTF, detection parameters) between sessions. Image stacks (pixel size ∼100 nm, z-step 1 μm) were acquired with 40x or 63x Plan Apo CS objectives with oil immersion.

### Image analysis

HABP and Ncan-GFP levels were quantified on 16-bit 40x (1.0 NA) confocal images acquired from coronal cortico-striatal organotypic slices. At least 3 slices were quantified per culture. The area of staining was calculated by manual thresholding in FIJI/ImageJ ^58^ and related to the total area of the image. To quantify HA network complexity by analysis of fractal dimension, HABP-Ncan-GFP double immunofluorescence images were segmented in both channels by thresholding, then binarized and skeletonized in FIJI. Analysis was restricted to areas with high AAV expression. Box counting fractal dimension was calculated with the FractalCount plugin for FIJI using default parameters. The length of hyaluronan cables was estimated on skeletonized images by quantifying the longest shortest path of the skeleton. Colocalization, fluorescence profiles, and maximal intensity projections were performed in FIJI. Images were linearly adjusted for brightness and contrast for visualization purposes, applying similar values to control-treated samples. 2-photon time-lapse recordings were registered with the StackReg plugin for FIJI, background was subtracted and a median filter of 1px was applied to eliminate small staining artifacts. In calcium transient imaging, images were processed with a Difference of Gaussians filter. Custom scripts for semi-automatization of the image analysis in FIJI are published in our GitHub repository (https://github.com/SoriaFN/Tools).

### *In vivo* two-photon microscopy

Two-photon microscopy was performed in 3–5-month-old C57BL/6J wild-type mice. A round craniotomy (around 4 mm in diameter) was made over the somatosensory cortex. 250 nl of AAV-Ncan-GFP 10^12^ vg/ml and 250 nl of AAV9-Syn-NES-jRGECO1a 10^10^ vg/ml were co-injected into the cortex at coordinate -0.3 dorsoventral at a rate of 60 nl/min using a motorized syringe pump. The craniotomy was covered with a glass coverslip (#1 thickness, diameter 4 mm) and an aluminum head holder was sealed using glue and dental cement. 30 days after the surgery, mice were anesthetized with isoflurane and placed on a heated pad, with the head fixed under a 20x (1.0 NA) objective (Olympus) in a Femto-2D microscope equipped with a tunable Ti::Sapphire MaiTai DeepSee femtosecond laser (Spectra Physics) and fluorescence filters to separate the Ncan-GFP signal (520/60) from jRGECO1 (605/70). Laser wavelength was tuned at 920 nm for Ncan-GFP and 1040 nm for jRGECO1. Laser power was 30 mW at the objective. Using blood vessels as ladmarks, images were taken in consecutive days in the same area. For imaging Ca^2+^ transients, isoflurane anesthesia was reduced at 0.5% to increase spontaneous neuronal activity.

### Statistics

Statistical analyses were performed using GraphPad Prism 8 (GraphPad Software Inc). Datasets were initially tested for normal distribution with D’Agostino-Pearson normality test. Normal populations were analyzed by unpaired two-tailed t-test (paired for HABP-Ncan-GFP co-localization comparisons). Continuous datasets (i.e. with a large number of data points) with non-normal distributions were analyzed by Kolmogorov–Smirnov (KS) test comparing cumulative distributions. For all statistical tests, the level of significance was set to P < 0.05. Exact P values are reported in each figure. In KS tests for large datasets, an approximate P value was calculated. Data are presented as mean ± standard error of mean (SEM) and *n* represents the number of animals or cultures analyzed (specified in figure legends). Continuous datasets are represented as relative frequency or cumulative distributions.

## Notes

### Competing Interest Statement

The authors have declared no competing interest.

